# PERK Signaling Regulates Extracellular Proteostasis of an Amyloidogenic Protein During Endoplasmic Reticulum Stress

**DOI:** 10.1101/419564

**Authors:** Isabelle C. Romine, R. Luke Wiseman

## Abstract

The PERK arm of the unfolded protein response (UPR) regulates cellular proteostasis and survival in response to endoplasmic reticulum (ER) stress. However, the impact of PERK signaling on extracellular proteostasis is poorly understood. We define how PERK signaling influences extracellular proteostasis during ER stress using a conformational reporter of the secreted amyloidogenic protein transthyretin (TTR). We show that inhibiting PERK signaling impairs ER stress-dependent secretion of destabilized TTR by increasing its ER retention in chaperone-bound complexes. Interestingly, PERK inhibition promotes the ER stress-dependent secretion of TTR in non-native conformations that accumulate extracellularly as soluble oligomers. Pharmacologic or genetic TTR stabilization partially restores secretion of native TTR tetramers. However, PERK inhibition still increases the ER stress-dependent secretion of TTR in non-native conformations under these conditions, indicating that the conformation of stable secreted proteins can also be affected by inhibiting PERK. Our results define a role for PERK in regulating extracellular proteostasis during ER stress and indicate that genetic or aging-related alterations in PERK signaling can exacerbate ER stress-related imbalances in extracellular proteostasis implicated in diverse diseases.

## INTRODUCTION

The PERK signaling arm of the unfolded protein response (UPR) has a critical role in defining cellular survival in response to pathologic insults that disrupt endoplasmic reticulum (ER) proteostasis (i.e., ER stress). PERK is activated in response to ER stress through a mechanism involving PERK dimerization and autophosphorylation [1, 2]. Once activated, PERK phosphorylates the α subunit of eukaryotic initiation factor 2 (eIF2α). This results in both a transient attenuation in new protein synthesis and the activation of stress-responsive transcription factors such as ATF4 [3-5]. PERK-dependent ATF4 activation induces expression of stress-responsive genes involved in diverse biologic functions including cellular redox, amino acid biosynthesis, and apoptotic signaling [3, 6, 7]. Apart from eIF2α, PERK also has been shown to activate NRF2 to promote cellular redox regulation during ER stress [8, 9]. Through this integration of transcriptional and translational signaling, PERK has a central role in dictating cellular proteostasis and survival in response to varying levels of ER stress. During acute ER insults, PERK signaling is important for regulating protective biologic functions including metabolite homeostasis, cellular redox homeostasis and mitochondrial function [8, 10-12]. However, chronic PERK activation caused by severe or persistent ER stress promotes apoptotic signaling primarily through the PERK-dependent transcriptional regulation of pro-apoptotic factors [13, 14].

Consistent with a role for PERK in dictating both protective and pro-apoptotic responses to specific ER insults, imbalances in PERK activity caused by genetic, environmental, or aging-related factors is implicated in the pathogenesis of diverse diseases. Sustained PERK signaling associated with chronic or severe ER insults is implicated in neurodegeneration associated with diseases such as Alzheimer’s disease and prion disease, [15, 16]. As such, pharmacologic inhibition of PERK has emerged as a potential strategy to ameliorate neurodegeneration-associated pathologies involved in these disorders [15]. In contrast, genetic and pharmacologic evidence demonstrates that reductions in PERK signaling can also influence disease pathogenesis. Loss-of-function mutations in *PERK* promote neonatal diabetes in mouse models and the human disease Wolcott-Rallison syndrome [17, 18]. Similarly, hypomorphic *PERK* alleles are implicated in the tau-associated neurodegenerative disorder progressive supranuclear palsy (PSP), suggesting that reduced PERK signaling promotes toxic tau aggregation [19, 20]. Consistent with this, pharmacologic PERK activation attenuates aggregation and toxicity of PSP-related tau mutants in mouse models [21]. Pharmacologic or chemical genetic increases in PERK signaling also reduce the toxic aggregation of rhodopsin mutants associated with retinal degeneration [19, 22, 23]. Thus, while significant focus has been directed to the pathologic importance of overactive or chronic PERK signaling in disease, it is clear that deficiencies in PERK activity also promote pathogenesis, reflecting a protective role for this UPR signaling arm in regulating cellular physiology in response to ER stress.

Interestingly, recent work has revealed PERK as a critical regulator of proteostasis within the ER – the first organelle of the secretory pathway. PERK-dependent translation attenuation regulates ER protein folding load in response to acute ER stress, freeing ER proteostasis factors to protect the secretory proteome from misfolding during the initial stages of toxic insult [2, 24]. In addition, PERK regulates both ER-to-Golgi anterograde trafficking and ER-associated degradation [25, 26], the latter a primary mechanism by which cells degrade ER proteins [27]. As such, genetic or pharmacologic inhibition of PERK signaling disrupts secretory proteostasis to reduce the secretion and increase the intracellular accumulation of proteins including collagen, insulin, or rhodopsin in high molecular weight (HMW) aggregates [28-31]. These results define an important role for PERK in regulating ER proteostasis in response to pathologic insults. However, considering the importance of the ER in regulating proteostasis in downstream secretory environments such as the extracellular space, a critical question remains: ‘‘*How does PERK signaling impact extracellular proteostasis in response to ER stress*?’.

Imbalances in ER proteostasis can propagate to extracellular environments through the secretion of proteins in non-native conformations that accumulate as toxic oligomers and aggregates associated with proteotoxicity in etiologically-diverse protein aggregation diseases, including many amyloid diseases [32-34]. Thus, impaired secretory proteostasis afforded by genetic or aging-related imbalances in PERK activity could exacerbate ER stress-dependent increases in the secretion of non-native proteins, challenging the conformational integrity of the secreted proteome and increasing extracellular populations of toxic oligomers. Here, we define how pharmacologic inhibition of PERK signaling impacts extracellular proteostasis during ER stress using a conformational reporter of the model secreted amyloidogenic protein transthyretin (TTR) – a natively tetrameric protein that aggregates into toxic oligomers and amyloid fibrils associated with diverse TTR-related amyloid diseases [35, 36]. We show that blocking PERK activity during ER stress exacerbates the stress-dependent secretion of both destabilized and wild-type TTR in non-native conformations that accumulate extracellularly as soluble oligomers commonly associated with TTR proteotoxicity [35]. These results demonstrate that alterations in PERK signaling can directly challenge extracellular proteostasis by reducing the secretion of native, functional proteins and promoting the extracellular accumulation of soluble oligomers associated with the pathogenesis of diverse types of protein aggregation diseases.

## RESULTS & DISCUSSION

### PERK inhibition disrupts secretion of destabilized TTR^A25T^ during ER stress

The inhibition of PERK signaling disrupts the trafficking of proteins such as collagen and insulin through the secretory pathway [28-31]. Here, we define how pharmacologic inhibition of PERK signaling influences secretion of the amyloidogenic protein transthyretin (TTR) during ER stress. Previously, we showed that administration of the ER stressor thapsigargin (Tg) reduces the secretion of destabilized, aggregation-prone TTR variants such as TTR^A25T^ [33, 37, 38]. To define how PERK inhibition influences the Tg-dependent reduction in TTR^A25T^ secretion, we pretreated HEK293T cells expressing FLAG-tagged TTR^A25T^ (^FT^TTR^A25T^) with Tg and/or the PERK inhibitor ISRIB for 16 h and then monitored ^FT^TTR^A25T^ secretion by [^35^S] metabolic labeling (**Fig. 1A**). ISRIB is an antagonist of PERK signaling that binds to eIF2B and desensitizes cells to eIF2α phosphorylation [39, 40]. Importantly, co-pretreatment with Tg and ISRIB increased metabolic labeling of ^FT^TTR^A25T^ relative to pretreatment with Tg alone (**Fig. 1A,B**). This is consistent with ISRIB blocking Tg-dependent translational attenuation downstream of PERK [39, 40]. Interestingly, co-pretreatment with ISRIB and Tg enhanced the Tg-dependent reduction in ^FT^TTR^A25T^ secretion (**Fig. 1A,C**). However, this enhanced reduction in ^FT^TTR^A25T^ secretion did not correspond with an increased loss of total ^FT^TTR^A25T^, indicating that co-pretreatment with Tg and ISRIB does not increase ^FT^TTR^A25T^ degradation (**Fig. 1A,D**). Instead, this pretreatment increased the accumulation of ^FT^TTR^A25T^ in lysate fractions, reflecting increased intracellular retention (**Fig. 1A,E**). Identical results were obtained using the PERK inhibitor GSK2656157 (herein referred to as GSK), which blocks PERK signaling by direct binding to the PERK kinase active site (**Fig. EV1A-E**)[41]. The use of the two mechanistically distinct inhibitors of PERK signaling indicates that the enhanced reduction in ^FT^TTR^A25T^ secretion observed in cells pretreated with Tg and PERK inhibitors cannot be attributed to off-pathway activities of these compounds [42]. Thus, these results show that pharmacologic inhibition of PERK signaling impairs secretory proteostasis for ^FT^TTR^A25T^ during ER stress by reducing its secretion and increasing its intracellular retention.

**Figure 1.**
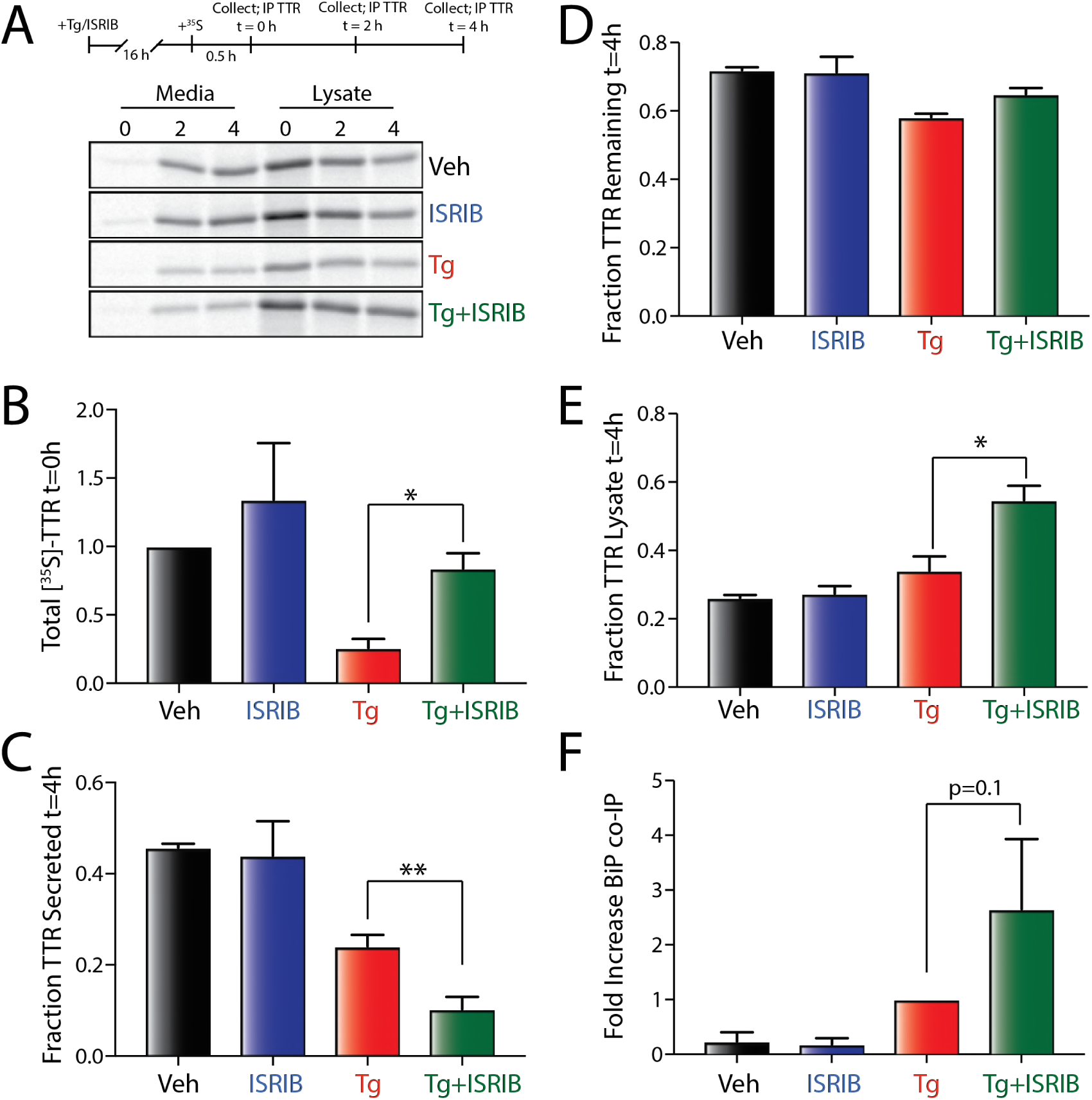
Pharmacologic PERK inhibition disrupts ^FT^TTR^A25T^ secretory proteostasis during ER stress. **A.** Representative autoradiogram of [^35^S]-labeled ^FT^TTR^A25T^ in lysates and media collected from HEK293T cells pretreated for 16 h with thapsigargin (Tg; 500 nM) and/or ISRIB (200 nM). The protocol for the experiment is shown above. **B.** Graph showing the total amount of [^35^S]-labeled ^FT^TTR^A25T^ at t = 0 h in HEK293T cells pretreated for 16 h with thapsigargin (Tg; 500 nM) and/or ISRIB (200 nM). Data are shown normalized to vehicle-treated cells. Error bars show SEM for n=3 independent experiments. A representative autoradiogram is shown in **Fig. 1A.** **C.** Graph showing the fraction [^35^S]-labeled ^FT^TTR^A25T^ secreted at t = 4 h in HEK293T cells pretreated for 16 h with thapsigargin (Tg; 500 nM) and/or ISRIB (200 nM). Fraction secreted was calculated as described in **Materials and Methods**. Error bars show SEM for n=3 independent experiments. A representative autoradiogram is shown in **Fig. 1A.** **D.** Graph showing the fraction [^35^S]-labeled ^FT^TTR^A25T^ remaining at t = 4 h in HEK293T cells pretreated for 16 h with thapsigargin (Tg; 500 nM) and/or ISRIB (200 nM). Fraction remaining was calculated as described in **Materials and Methods**. Error bars show SEM for n=3 independent experiments. A representative autoradiogram is shown in **Fig. 1A.** **E.** Graph showing the fraction [^35^S]-labeled ^FT^TTR^A25T^ in lysates at t = 4 h in HEK293T cells pretreated for 16 h with thapsigargin (Tg; 500 nM) and/or ISRIB (200 nM). Fraction lysate was calculated as described in **Materials and Methods**. Error bars show SEM for n=3 independent experiments. A representative autoradiogram is shown in **Fig. 1A.** **F.** Graph showing the recovery of BiP in anti-FLAG IPs from HEK293T cells expressing ^FT^TTR^A25T^ and pretreated for 16 h with Tg (500 nM) and/or ISRIB (200 nM). Data are shown normalized to Tg-treated cells. Error bars show SEM for n=3 independent experiments. A representative immunoblot is shown in **Fig. EV1F**. *indicates p<0.05; **indicates p<0.01 for a paired two-tailed t-test.

### PERK inhibition increases ER stress-dependent interactions between TTR^A25T^ and ER chaperones

The intracellular retention of ^FT^TTR^A25T^ observed upon co-pretreatment with Tg and ISRIB likely reflects increased accumulation of non-native TTR conformations within the ER, as observed for other proteins [28, 29, 31]. To test this, we measured the intracellular interaction between ^FT^TTR^A25T^ and ER chaperones, which bind non-native protein conformations, in HEK293T cells treated with Tg and/or ISRIB for 16 h using an established immunopurification (IP)/immunoblotting (IB) assay [38]. Treatment with Tg increases the relative recovery of the ER ATP-dependent HSP70 chaperone BiP and the BiP co-chaperones ERdj3 and HYOU1 in ^FT^TTR^A25T^ IPs (**Fig. 1F** and **Fig. EV1F**). This indicates that Tg treatment increases the non-native population of intracellular TTR that can engage ER chaperones. Co-treatment with Tg and ISRIB further increased the relative interactions between ^FT^TTR^A25T^ and these ER chaperones, as compared to Tg treatment alone. Importantly, we do not observe significant differences in the intracellular levels of BiP or HYOU1 under these conditions (**Fig. EV1F**). This indicates that ISRIB-dependent inhibition of PERK signaling does not globally disrupt ER stress-dependent upregulation of these UPR target genes. This suggests that disruption of ^FT^TTR^A25T^ secretion afforded by PERK inhibition likely results from translational attenuation, and not impaired transcriptional regulation of these ER chaperones. However, we cannot rule out the possibility that impaired transcription of other UPR-regulated proteins contribute to the altered secretion of ^FT^TTR^A25T^ observed in cells co-treated with Tg and ISRIB. Collectively, these results indicate that the increased accumulation of destabilized ^FT^TTR^A25T^ afforded by co-treatment with PERK inhibitors and ER stress reflects increased ER retention of non-native TTR in chaperone-bound complexes. This suggests that PERK inhibition during ER stress disrupts the ability of cells to properly fold and assemble ^FT^TTR^A25T^ into its native tetrameric conformation.

### PERK inhibition reduces the population of ^FT^TTR^A25T^ secreted as native tetramers

Imbalances in ER proteostasis can propagate to extracellular environments through the secretion of proteins such as TTR in non-native conformations [32, 33]. Thus, we used a previously established assay to define how the ER accumulation of non-native ^FT^TTR^A25T^ observed upon co-treatment with PERK inhibitors and ER stress impacts the conformation of ^FT^TTR^A25T^ secreted from mammalian cells (**Fig. 2A**)[33]. In this assay, we collect conditioned media from cells expressing ^FT^TTR^A25T^ prepared in the presence of the cell impermeable compound tafamidis-sulfonate (Taf-S; **Fig. EV2A**) – a compound that immediately binds and stabilizes native TTR tetramers once secreted to the extracellular space [33]. We then quantify total, tetrameric, and aggregate TTR in this conditioned media. Total TTR is quantified by SDS-PAGE/IB. Tetrameric TTR is measured by incubating conditioned media with compound **1** (**Fig. EV2A**) – a fluorogenic compound that readily displaces the Taf-S in the two small-molecule binding sites of the native TTR tetramer and becomes fluorescent upon binding [33, 43, 44]. Tetramers are then separated by anion exchange chromatography and quantified by compound **1** fluorescence (**Fig. EV2B**). Finally, soluble TTR aggregates are monitored by Clear Native (CN)- PAGE/IB [33, 34]. We previously used this approach to show that Tg-induced ER stress reduces the secretion of destabilized ^FT^TTR^A25T^ as native tetramers and increases accumulation of this destabilized, aggregation-prone protein as soluble oligomers in conditioned media [33].

**Figure 2.**
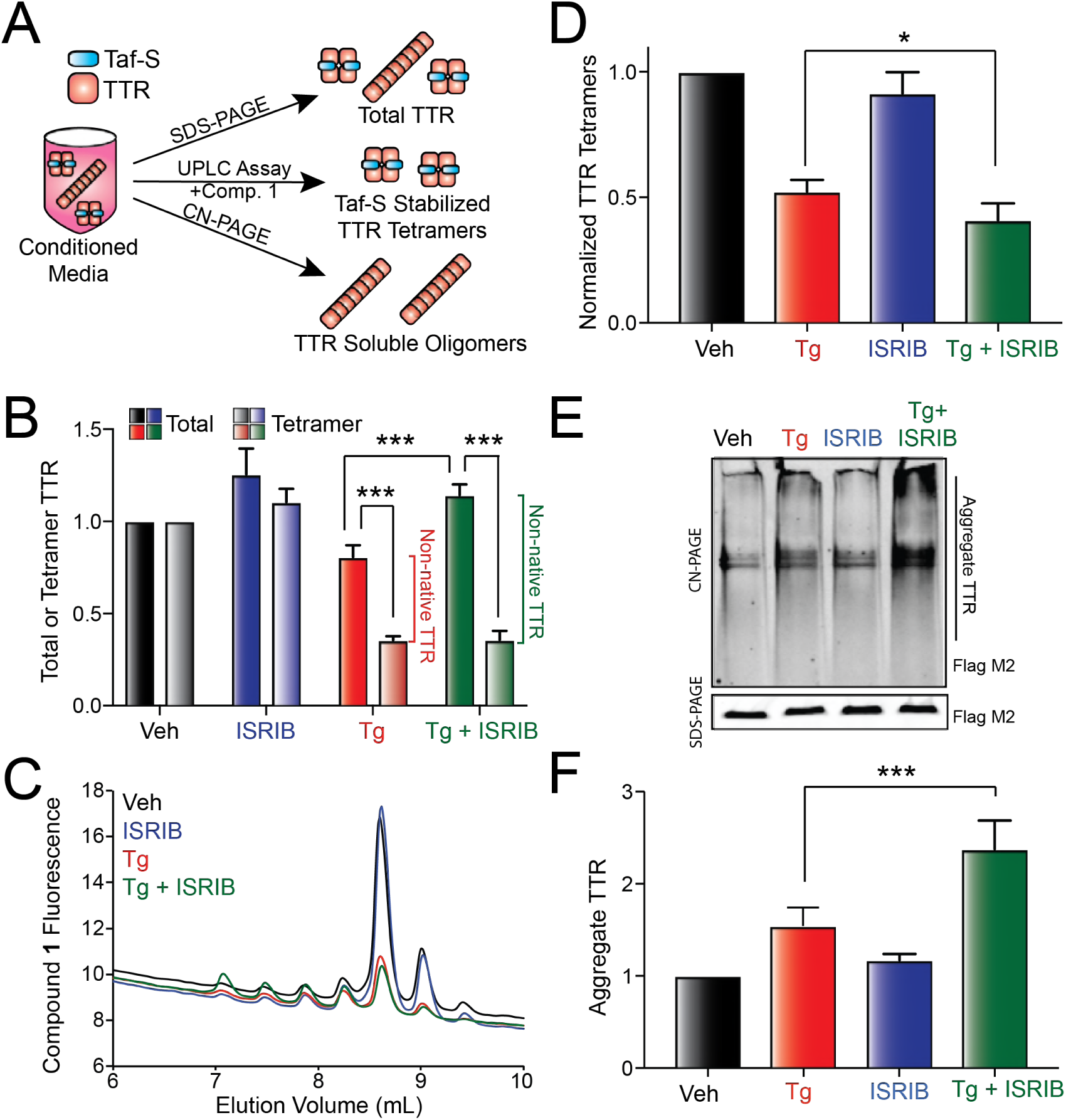
Pharmacologic inhibition of PERK signaling increases secretion of ^FT^TTR^A25T^ in non-native conformations. **A.** Illustration showing the three assays used to monitor total, tetrameric, and aggregate TTR in conditioned media. **B.** Graph showing total (dark colors) and tetrameric (light colors) ^FT^TTR^A25T^ in conditioned media prepared in the presence of Taf-S on HEK293T cells treated for 16 h with Tg (500 nM) and/or ISRIB (200 nM). Total and tetrameric ^FT^TTR^A25T^ are normalized to vehicle. Relative populations of non-native TTR in each condition are shown. Error bars show SEM for n=13 independent experiments. **C.** Representative chromatogram showing ^FT^TTR^A25T^ tetramers in conditioned media prepared in the presence of Taf-S on HEK293T cells treated for 16 h with Tg (500 nM) and/or ISRIB (200 nM). ^FT^TTR^A25T^ tetramers were separated on anion exchange chromatography using the UPLC system and visualized by compound **1** fluorescence. **D.** Graph showing normalized ^FT^TTR^A25T^ tetramers in conditioned media prepared in the presence of Taf-S on HEK293T cells treated for 16 h with Tg (500 nM) and/or ISRIB (200 nM). Normalized TTR tetramers were calculated using the formula: normalized TTR tetramers for a given condition = (TTR tetramer signal condition / TTR tetramer signal veh) / (total TTR signal condition / TTR tetramer signal veh). Error bars show SEM for n=13 independent experiments. **E.** Representative Clear Native (CN)-PAGE immunoblot of ^FT^TTR^A25T^ soluble aggregates in conditioned media prepared in the presence of Taf-S on HEK293T cells treated for 16 h with Tg (500 nM) and/or ISRIB (200 nM). Total ^FT^TTR^A25T^ in each condition is shown in the SDS-PAGE immunoblot. **F.** Graph showing the amounts of ^FT^TTR^A25T^ soluble aggregates in conditioned media prepared in the presence of Taf-S on HEK293T cells treated for 16 h with Tg (500 nM) and/or ISRIB (200 nM). Aggregate ^FT^TTR^A25T^ is normalized to vehicle. Error bars show SEM for n=13 independent experiments. *indicates p<0.05, ***indicates p<0.005 for a two-tailed paired t-test.

To define how PERK inhibition influences ER stress-dependent alterations in the conformation of secreted ^FT^TTR^A25T^, we conditioned media in the presence of Taf-S for 16 h on HEK293T cells expressing ^FT^TTR^A25T^ and treated with Tg and/or ISRIB. We then measured total, tetrameric, and aggregate TTR using the assays described above (**Fig. 2A**). Treatment with Tg reduced the accumulation of total ^FT^TTR^A25T^ in conditioned media, consistent with published results (**Fig. 2B**)[33, 38]. However, despite pretreatment with Tg and ISRIB reducing the fraction ^FT^TTR^A25T^ secreted (**Fig. 1C**), the accumulation of ^FT^TTR^A25T^ in conditioned media over the 16 h incubation was not reduced relative to vehicle-treated cells (**Fig. 2B**). This likely reflects the inhibition of PERK-regulated translational attenuation afforded by ISRIB over the duration of the media conditioning (**Fig. 1B**).

We next monitored tetrameric ^FT^TTR^A25T^ in this conditioned media using our compound **1** fluorescence/anion exchange chromatography assay (**Fig. 2A**). As reported previously [33], ^FT^TTR^A25T^ tetramers in conditioned media migrate as multiple peaks by anion exchange chromatography, reflecting the distribution of heterotetramers consisting of unmodified subunits and subunits containing posttranslational modifications such as sulfonylation and cysteinylation (**Fig. 2C**). Co-treatment with Tg and ISRIB significantly altered the distribution of these peaks, resulting in an increase of heterotetramers eluting earlier on the anion exchange column (**Fig. 2C**, green). This suggests that Tg and ISRIB co-treatment alters posttranslational modifications of ^FT^TTR^A25T^ secreted to the extracellular space. As expected, liquid chromatography (LC)-mass spectrometry (MS) analysis of ^FT^TTR^A25T^ IP’d from conditioned media prepared on cells co-treated with Tg and ISRIB shows increased accumulation of unmodified ^FT^TTR^A25T^ (**Fig. EV2C**). This is consistent with the earlier migration of tetramers observed under these conditions by anion exchange chromatography and demonstrates that PERK inhibition disrupts the secretory proteostasis environment to influence posttranslational TTR modifications during ER stress.

To quantify the recovery of ^FT^TTR^A25T^ tetramers in conditioned media, we integrated the fluorescence across all of the tetramer peaks. We observed significant reductions in the recovery of ^FT^TTR^A25T^ tetramers in media prepared on Tg-treated cells (**Fig. 2B**), consistent with published results [33]. However, despite lower levels of total ^FT^TTR^A25T^ in media conditioned on cells treated with Tg relative to cells treated with Tg and ISRIB, identical amounts of tetramers were detected in conditioned media prepared under these two conditions (**Fig. 2B**). This indicates that PERK inhibition decreases the relative population of ^FT^TTR^A25T^ secreted as native tetramers. This can be demonstrated by normalizing the recovered ^FT^TTR^A25T^ tetramers by the total amount of ^FT^TTR^A25T^ in media prepared on cells treated with Tg and/or ISRIB [33]. This normalization confirms that PERK inhibition reduces the population of ^FT^TTR^A25T^ secreted as native tetramers during ER stress (**Fig. 2D**). Thus, our results show that PERK inhibition increases the ER-stress-dependent secretion of ^FT^TTR^A25T^ in non-native conformations.

### Inhibiting PERK increases the extracellular accumulation of ^FT^TTR^A25T^ oligomers during ER stress

Non-native ^FT^TTR^A25T^ accumulates as soluble oligomers in conditioned media owing to the low stability of the aggregation-prone ^FT^TTR^A25T^ variant [33, 34]. Thus, the increased *total* secreted TTR (**Fig. 2B**) combined with the increased secretion of ^FT^TTR^A25T^ in non-native conformations (**Fig. 2D**) in cells co-treated with Tg and ISRIB should be reflected by increased accumulation of extracellular soluble oligomers in media prepared under these conditions. Tg treatment alone significantly increased the accumulation of soluble ^FT^TTR^A25T^ aggregates (**Fig. 2E,F**). This is consistent with previous results and further demonstrates the ER stress-dependent increase in the secretion of non-native ^FT^TTR^A25T^ [33]. Interestingly, co-treatment with Tg and ISRIB significantly increased the accumulation of ^FT^TTR^A25T^ soluble aggregates in conditioned media relative to treatment with Tg alone (**Fig. 2E,F**), consistent with the increase in non-native ^FT^TTR^A25T^ secreted under these conditions (**Fig. 2B,D**). These results demonstrate that inhibiting PERK signaling during ER stress directly challenges extracellular proteostasis by exacerbating the ER stress-dependent secretion of destabilized ^FT^TTR^A25T^ in non-native conformations that accumulate extracellularly as soluble oligomers.

### Pharmacologic stabilization of TTR tetramers partially rescues the secretion of native ^FT^TTR^A25T^ tetramers in cells treated with PERK inhibitors and ER stress

TTR tetramers can be stabilized intracellularly by administration of the cell permeable kinetic stabilizer tafamidis (Taf) [33, 37]. Taf-dependent intracellular stabilization prevents the dissociation of tetramers prior to secretion to the extracellular space. Thus, conditioning media in the presence of Taf provides an opportunity to determine whether the increased populations of ^FT^TTR^A25T^ secreted from cells co-treated with Tg and ISRIB reflects increased dissociation of tetramers during the secretion process (i.e., reduced tetramer stability) and/or increased secretion of ^FT^TTR^A25T^ prior to tetramer assembly (i.e., reduced tetramer assembly). To test this, we conditioned media on HEK293T cells expressing ^FT^TTR^A25T^ treated with Tg and/or ISRIB in the presence of either Taf-S or Taf. We then monitored total, tetrameric, and aggregate TTR using the assays shown in **Fig. 2A**.

Conditioning media in the presence of Taf significantly increased the recovery of ^FT^TTR^A25T^ tetramers in every condition, relative to media conditioned in the presence of Taf-S (**Fig. 3A,B** and **Fig. EV3**). This indicates that Taf-dependent intracellular stabilization of ^FT^TTR^A25T^ tetramers increases secretion of ^FT^TTR^A25T^ as native tetramers. This increase cannot be attributed to alterations in ^FT^TTR^A25T^ posttranslational modifications, as the addition of Taf does not alter the shift in tetramer distribution observed in cells co-treated with Tg and ISRIB (**Fig. 3B**). Importantly, the increased secretion of ^FT^TTR^A25T^ tetramers corresponds with reductions in the accumulation of soluble ^FT^TTR^A25T^ aggregates in conditioned media (**Fig. 3C**). This demonstrates that Taf-dependent intracellular stabilization of TTR tetramers reduces the secretion of ^FT^TTR^A25T^ in non-native conformations under all conditions. This indicates that the accumulation of non-native TTR observed under these conditions partially reflects dissociation of TTR tetramers (that can be stabilized intracellularly by Taf) within the secretory pathway. This reveals a specific advantage for employing kinetic stabilizing compound such as Taf that are cell permeable, as these compounds can stabilize aggregation-prone proteins intracellularly and mitigate potential imbalances in secretory proteostasis associated with the aberrant secretion of non-native conformations.

**Figure 3.**
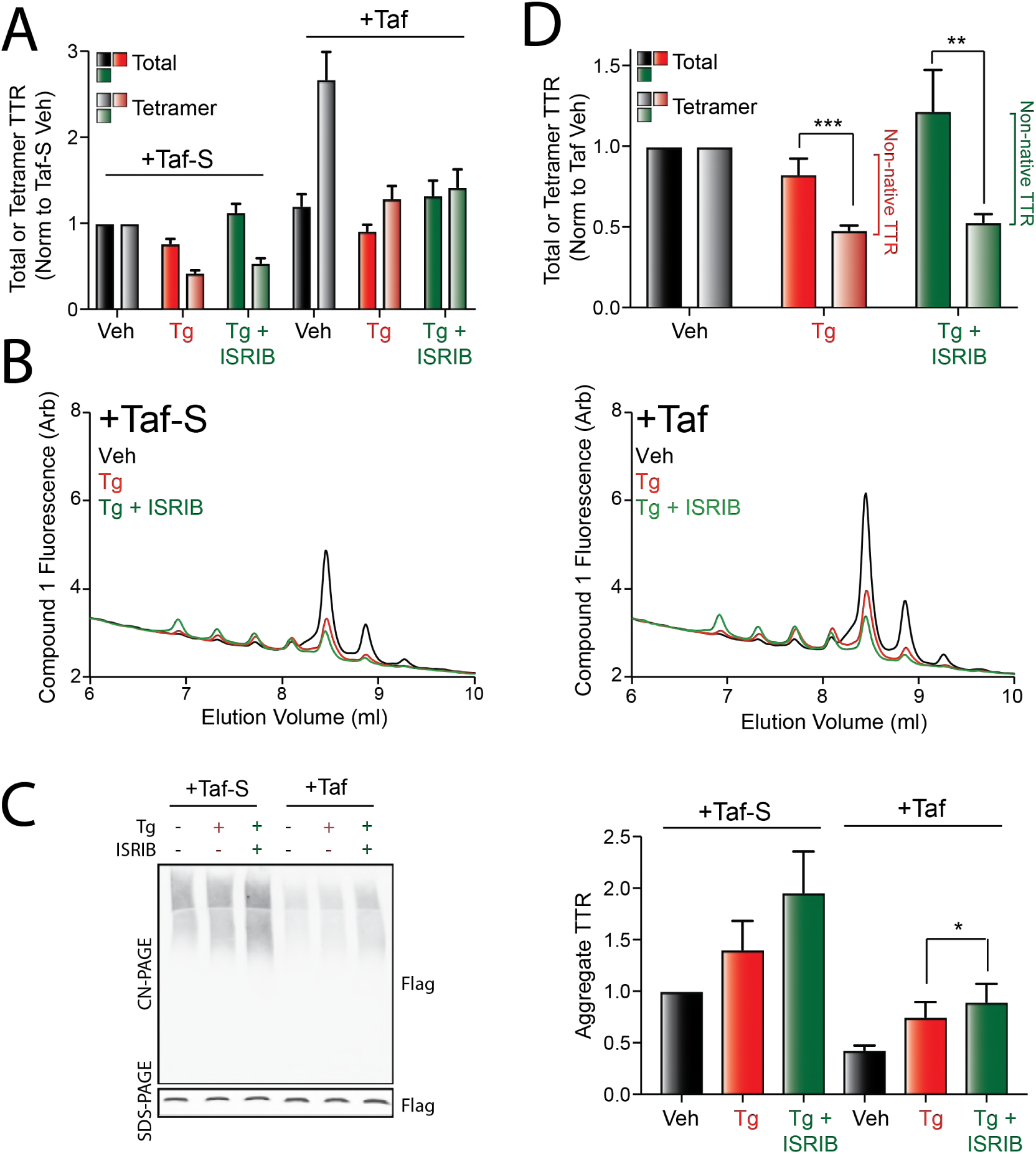
Pharmacologic stabilization of ^FT^TTR^A25T^ tetramers partially restores secretion in native conformations. **A.** Graph showing total (dark colors) and tetrameric (light colors) ^FT^TTR^A25T^ in conditioned media prepared in the presence of Taf-S or Taf on HEK293T cells treated for 16 h with Tg (500 nM) and/or ISRIB (200 nM). Total and tetrameric ^FT^TTR^A25T^ are normalized to media prepared on vehicle-treated cells in the presence of Taf-S. Error bars show SEM for n=8 independent experiments. **B.** Representative chromatogram showing ^FT^TTR^A25T^ tetramers in conditioned media prepared in the presence of Taf-S (left) or Taf (right) on HEK293T cells treated for 16 h with Tg (500 nM) and/or ISRIB (200 nM). ^FT^TTR^A25T^ tetramers were separated on anion exchange chromatography using the UPLC system and visualized by compound **1** fluorescence. **C.** Representative CN-PAGE/immunoblot (left) and quantification (right) of ^FT^TTR^A25T^ aggregates in conditioned media prepared in the presence of Taf-S or Taf on HEK293T cells treated for 16 h with Tg (500 nM) and/or ISRIB (200 nM). Total ^FT^TTR^A25T^ in each condition is shown in the SDS-PAGE immunoblot. **D.** Error bars show SEM for n=5 independent experiments. **E.** Graph showing total (dark colors) and tetrameric (light colors) ^FT^TTR^A25T^ in conditioned media prepared in the presence of Taf-S or Taf on HEK293T cells treated for 16 h with Tg (500 nM) and/or ISRIB (200 nM). Total and tetrameric ^FT^TTR^A25T^ are normalized to media prepared on vehicle-treated cells in the presence of Taf. Relative populations of non-native TTR in each condition are shown. Error bars show SEM for n=8 independent experiments. *indicates p<0.05, **p<0.01, and ***p<0.005 for a two-tailed paired t-test.

Despite the increased secretion of ^FT^TTR^A25T^ tetramers afforded by Taf-dependent intracellular stabilization, co-administration of Tg and ISRIB in the presence of Taf still increased secretion of ^FT^TTR^A25T^ in non-native conformations, relative to Tg treatment alone (**Fig. 3A,D**). Consistent with this, conditioned media prepared on cells co-treated with Tg and ISRIB in the presence of Taf show higher levels of soluble ^FT^TTR^A25T^ oligomers relative to Tg-treated cells, albeit significantly less than that observed for media conditioned in the presence of Taf-S (**Fig. 3C**). This demonstrates that despite the ability for Taf to stabilize a large population of ^FT^TTR^A25T^ tetramers in the secretory environment, PERK inhibition can still increase in the secretion of ^FT^TTR^A25T^ in non-native conformations under these conditions.

### Secretion of stable, wild-type TTR as native tetramers is reduced in cells co-treated with ER stress and PERK inhibition

The ability for PERK inhibition to promote ER stress-dependent secretion of ^FT^TTR^A25T^ in non-native conformations even in the presence of the stabilizing ligand Taf suggests that stable, wild-type TTR (TTR^WT^) could similarly be susceptible to reductions in tetramer secretion under these conditions. Initially, we confirmed that PERK inhibition disrupts secretory proteostasis for stable ^FT^TTR^WT^ by monitoring the interactions between ^FT^TTR^WT^ and ER chaperones in HEK293T cells treated with Tg and/or the PERK inhibitor GSK. Co-treatment with Tg and GSK significantly increased relative interactions between ^FT^TTR^WT^ and the ER chaperones BiP, ERdj3, and HYOU1 (**Fig. 4A**), mimicking the results observed with destabilized ^FT^TTR^A25T^ (**Fig. 1F** and **Fig. EV1F**). Furthermore, co-treatment with Tg and GSK increased intracellular levels of ^FT^TTR^WT^, suggesting that this condition leads to intracellular accumulation of stable ^FT^TTR^WT^ (**Fig. EV4A**). However, we did not observe significant increases in intracellular levels of BiP or HYOU1, further demonstrating that PERK inhibition does not impact Tg-dependent increases in these chaperones (**Fig. EV4A**). Collectively, these results indicate that PERK inhibition during ER stress disrupts secretory proteostasis of stable, wild-type TTR by increasing its ER retention in chaperone-bound complexes – results analogous to that observed for destabilized ^FT^TTR^A25T^ (**Fig. 1F** and **Fig. EV1F**).

**Figure 4.**
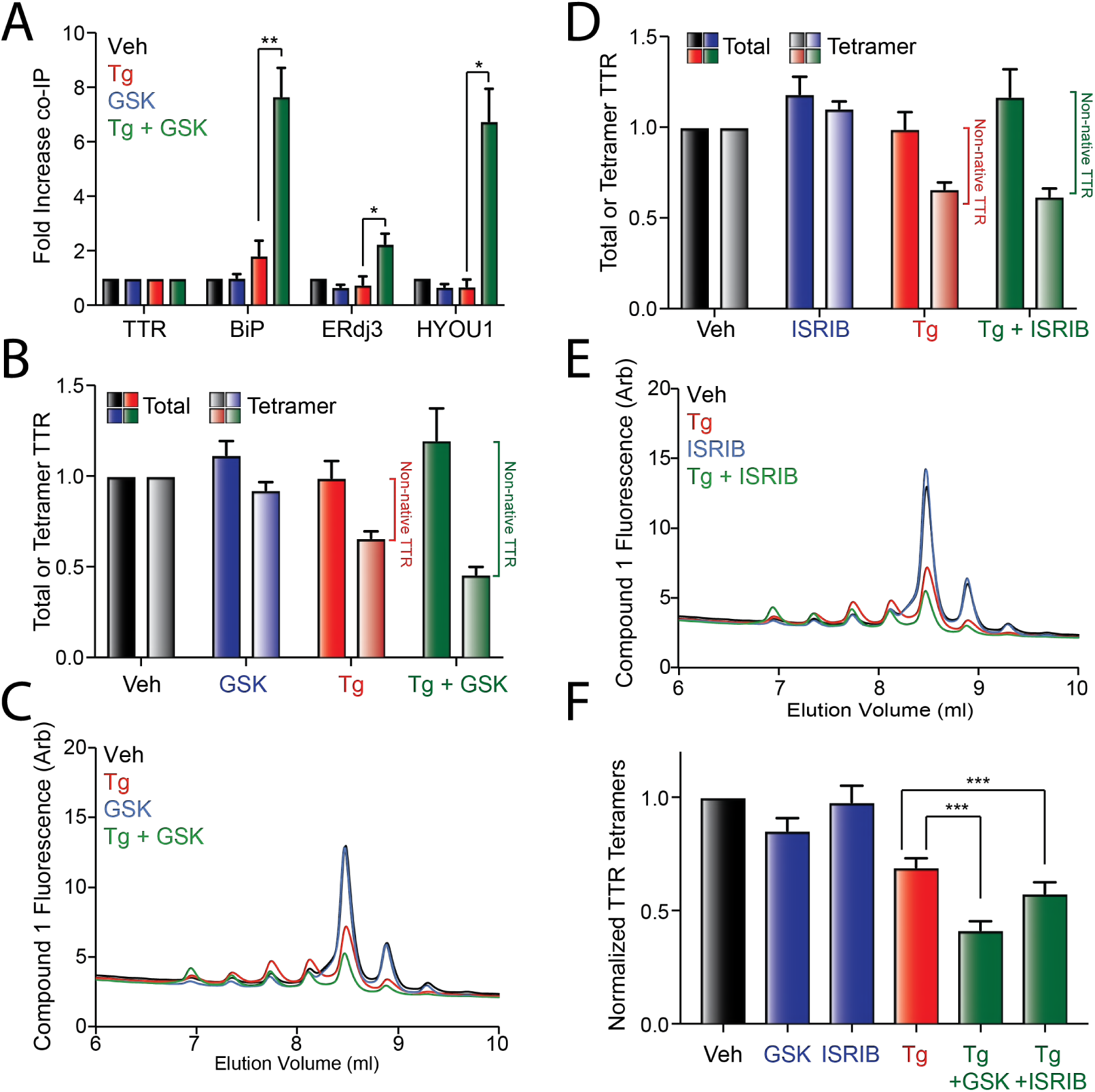
PERK inhibition promotes secretion of stable, wild-type TTR in non-native conformations during ER stress. **A.** Graph showing the recovery of TTR, BiP, ERdj3, and HYOU1 in anti-FLAG IPs from HEK293T cells expressing ^FT^TTR^WT^ and pretreated for 16 h with Tg (500 nM) and/or GSK (300 nM). Error bars show SEM for n=4 independent experiments. A representative immunoblot is shown in **Fig. EV4A.** **B.** Graph showing total (dark colors) and tetrameric (light colors) ^FT^TTR^WT^ in conditioned media prepared in the presence of Taf-S on HEK293T cells treated for 16 h with Tg (500 nM) and/or GSK (300 nM). Total and tetrameric ^FT^TTR^WT^ are normalized to media prepared on vehicle-treated cells. Relative populations of non-native TTR in each condition are shown. Error bars show SEM for n=10 independent experiments. **C.** Representative chromatogram showing ^FT^TTR^WT^ tetramers in conditioned media prepared in the presence of Taf-S on HEK293T cells treated for 16 h with Tg (500 nM) and/or GSK (300 nM). ^FT^TTR^WT^ tetramers were separated on anion exchange chromatography using the UPLC system and visualized by compound **1** fluorescence. **D.** Graph showing total (dark colors) and tetrameric (light colors) ^FT^TTR^WT^ in conditioned media prepared in the presence of Taf-S on HEK293T cells treated for 16 h with Tg (500 nM) and/or ISRIB (200 nM). Total and tetrameric ^FT^TTR^WT^ are normalized to media prepared on vehicle-treated cells. Relative populations of non-native TTR in each condition are shown. Error bars show SEM for n=10 independent experiments. **E.** Representative chromatogram showing ^FT^TTR^WT^ tetramers in conditioned media prepared in the presence of Taf-S on HEK293T cells treated for 16 h with Tg (500 nM) and/or ISRIB (200 nM). ^FT^TTR^WT^ tetramers were separated on anion exchange chromatography using the UPLC system and visualized by compound **1** fluorescence. **H.** Graph showing normalized ^FT^TTR^WT^ tetramers in conditioned media prepared in the presence of Taf-S on HEK293T cells treated for 16 h with Tg (500 nM), ISRIB (200 nM), and/or GSK (300 nM). Normalized TTR tetramers were calculated using the formula: normalized TTR tetramers for a given condition = (TTR tetramer signal condition / TTR tetramer signal veh) / (total TTR signal condition / TTR tetramer signal veh). Error bars show SEM for n=10. p<0.05; p<0.01; p<0.005 for two-tailed paired t-test.

Next, we conditioned media in the presence of Taf-S on HEK293T cells expressing ^FT^TTR^WT^ treated with Tg and/or GSK and monitored the recovery of total, tetrameric, or aggregate TTR using the assays described in **Fig. 2A**. Tg and GSK co-treatment increased the population of ^FT^TTR^WT^ secreted in non-native conformations (**Fig. 4B,C** and **Fig. EV4B**). Identical results were observed in cells co-treated with Tg and ISRIB (**Fig. 4D,E**). Normalizing the recovered ^FT^TTR^WT^ tetramers by the total amount of ^FT^TTR^WT^ in conditioned media confirms that PERK inhibition reduces the population of stable ^FT^TTR^WT^ secreted as tetramers during ER stress (**Fig. 4F**). Unfortunately, the stability of ^FT^TTR^WT^ monomers (relative to ^FT^TTR^A25T^ monomers) combined with the relatively low concentrations of non-native ^FT^TTR^WT^ in conditioned media hinders the aggregation of this protein in the extracellular space, making it difficult to reproducibly visualize ^FT^TTR^WT^ soluble aggregates in conditioned media [45]. However, co-treatment with Tg and GSK does appear to increase the population of ^FT^TTR^WT^ soluble aggregates observed by CN-PAGE/IB, consistent with the increased secretion of this protein in non-native conformations (**Fig. EV4C**). Collectively, these results show that PERK inhibition increases the ER stress-dependent secretion of wild-type TTR in non-native conformations, indicating that PERK inhibition can disrupt extracellular proteostasis of stable secreted proteins.

### Concluding Remarks

Imbalances in PERK signaling are implicated in the onset and pathogenesis of etiologically-diverse diseases [15-23]. Here, we show that PERK inhibition exacerbates the ER stress-dependent secretion of the disease relevant amyloidogenic protein TTR in non-native conformations that accumulate extracellularly as soluble oligomers commonly associated with TTR proteotoxicity [35]. This indicates that PERK signaling has an important role in dictating extracellular proteostasis by controlling the conformational integrity of secreted proteins. As such, genetic, aging-related, or pharmacologic conditions that reduce PERK signaling can lead to pathologic imbalances in extracellular proteostasis that contributes to human disease. The increased secretion of proteins in non-native conformations will increase extracellular populations available for concentration-dependent aggregation into toxic oligomers and amyloid fibrils, as we show for TTR. Alternatively, reduced secretion of proteins in native conformations can also prevent important functions of secreted proteins in biologic activities such as hormonal signaling, cell-cell communication and immunological signaling. Thus, our demonstration that reduction in PERK activity can impact the conformation of secreted proteins such as TTR reveals a new mechanism that can contribute to the pathogenesis of diseases associated with genetic or aging-related reductions in PERK activity. Furthermore, our results reveal a new consideration when employing PERK inhibitors to attenuate chronic PERK activation in disease, as this approach can promote imbalances in extracellular proteostasis and function that could be detrimental for long-term organismal survival

## MATERIALS AND METHODS

### Plasmids, Antibodies and Reagents

The ^FT^TTR^A25T^ and ^FT^TTR^WT^ plasmids were prepared in the pcDNAI vector as previously reported [38]. Antibodies were purchased from commercial sources: anti-ERdj3/DNAJB11 (Proteintech Group; Cat #15484-1-AP), anti-KDEL [10C3] (ENZO; Cat #ADI-SPA827-F), anti-HYOU1/ORP150 [C2C3] (Genetex; Cat#GTX102255), anti-TTR (Agilent; Cat #A000202-2), and anti-Flag M2 (Sigma Aldrich, Cat #F1804). Secondary antibodies for immunoblotting including IRDye Goat anti-mouse 800CW and IRDye Goat anti-rabbit 680CW were purchased from LICOR. Anti-FLAG immunopurifications were performed using anti-Flag M1 Agarose Affinity Gel (Sigma Aldrich; A4596). Tafamidis (Taf), Tafamidis-sulfonate (Taf-S), and compound **1** were generously gifted by Jeffery Kelly at TSRI. DMEM (Cat # 15-017-CM) was purchased from Cellgro. Penicillin/streptomycin (Cat# 15140122) glutamine (Cat# 25030081), and 0.25% Trypsin/EDTA (Cat# 25200056) were purchased from Invitrogen. Fetal Bovine Serum was purchased from Hyclone (Cat# 30396.03; Lot# AB10136011). Thapsigargin was purchased from Fisher Scientific (Cat #50-464-295). ISRIB (Cat #SML0843) was purchased from Sigma. GSK2656157 (Cat# 9466) was purchased from BioVision.

### Cell Culture, Plasmids, Transfections, and Lysates

HEK293T cells were cultured in DMEM supplemented 10% FBS, Penicillin/Streptomycin and glutamine. Cells were transfected with ^FT^TTR^A25T^/^FT^TTR^WT^ in pcDNAI using calcium phosphate transfection, as previously reported [38]. Cells were incubated in transfection media overnight and media was replaced with fresh supplemented complete media the next morning. Cells were lysed with 20 mM HEPES pH 7.2, 100 mM NaCl, 1% Triton X, 1 mM EDTA unless otherwise stated. Cell lysates were normalized to over all protein concentration in each sample using a Bradford assay.

### [^35^S] metabolic labeling assay

Metabolic labeling assays were performed as previously described [37, 38]. Briefly, HEK293T cells transfected with ^FT^TTR^A25T^ pretreated for 16 h with the indicated condition were labeled for 30 min with EasyTag Express ^35^S Protein Labeling Mix (0.1mCi/mL; Perkin Elmer) in DMEM lacking both Cys and Met (Invtrogen) supplemented with 10% dialyzed FBS, penicillin/streptomycin, and glutamine. Cells were then washed and incubated in complete media. At the indicated time, media and lysates were collected. ^FT^TTR^A25T^ was then immunopurified from the lysate and media using anti-Flag M1 Agarose Affinity Gel (Sigma Aldrich) and incubated overnight. We then washed the beads and eluted ^FT^TTR^A25T^ by boiling the beads in 3x Laemmli buffer including 100 mM DTT. Proteins were separated on a 12% SDs-PAGE gel, dried, exposed to phosphorimager plates (GE Healthcare), and imaged with a Typhoon imager. Band intensities were quantified by densitometry using ImageQuant. Fraction secreted was calculated using the equation: fraction secreted = [^35^S]-TTR signal in media at t / ([^35^S]-TTR signal in media at t=0h + [^35^S]-TTR signal in lysate at t=0h). Fraction remaining was calculated using the equation: fraction remaining = ([^35^S]-TTR signal in media at t + [^35^S]-TTR signal in lysate at t) / ([^35^S]-TTR signal in media at t=0h + [^35^S]-TTR signal in lysate at t=0h). Fraction lysate was calculated using the equation: fraction lysate = [^35^S]-TTR signal in lysate at t / ([^35^S]-TTR signal in media at t=0h + [^35^S]-TTR signal in lysate at t=0h).

### Media Conditioning, SDS-PAGE, CN-PAGE and Immunoblotting

HEK293T cells transfected with ^FT^TTR^A25T^/^FT^TTR^WT^ were incubated overnight. The following morning media was changed and cells were incubated for 1 hour. Cells were then replated into a poly-D-lysine coated 6-well plate. The next evening, cells were treated with the indicated treatment for 16 hours and conditioned in 1 mL of media. Conditioned media was then collected and cleared from large debris by centrifugation at 1000 x g for 10 min. For SDS-PAGE, samples were boiled in Lammeli Buffer containing 100 mM DTT for 5 minutes and resolved on a 12% acrylamide gel. Gels were transferred to 0.2 μm nitrocellulose membranes at 100V for 60 minutes. For CN-PAGE, samples were added to 5× Clear Native Page Buffer (final concentration: 0.1% Ponceau Red, 10% glycerol, 50mM 6-aminohexanoic acid, 10mM Bis-Tris pH 7.0) and resolved at 4°C on a Novex NativePage 4-16% Bis-Tris Protein Gel run at 150 V for 3 hours. Protein was transferred at 100 V for 90 minutes to a 0.2 μm nitrocellulose membrane. Membranes were incubated with the indicated primary antibody overnight and then incubated with the appropriate LICOR secondary antibody. Protein bands where then visualized and quantified using the LICOR Odyssey Infrared Imaging System.

### Quantification of ^FT^TTR tetramers in conditioned media

^FT^TTR tetramers in conditioned media were quantified as previously described [33]. Briefly, conditioned media was prepared on HEK293T cells expressing the indicated ^FT^TTR variant treated with the indicated conditions. Conditioned media was then incubated with 5 μM compound **1** overnight. Media samples (60 μL) were separated over a Waters Protein-Pak Hi Res Q, 5 μm 4.6 x 100 mm anion exchange column in 25 mm Tris pH 8.0, 1 mM EDTA with a linear 1M NaCl gradient using a ACQUITY UPLC H-Class Bio System (Waters). Fluorescence of TTR tetramers conjugated to compound **1** was observed by excitation at 328 nm and emission at 430 nm. Peaks were integrated and data was collected using Empower 3 software according to the manufacture’s protocol.

### Liquid chromatography (LC)-mass spectrometry of immunopurified ^FT^TTR

Media was conditioned for 18 h on HEK293T cells transfected with ^FT^TTR^A25T^ treated with the indicated condition. Media was then collected and cleared from debris by centrifugation at 1000 x for 10 min. ^FT^TTR^A25T^ was immunopurified from the conditioned media using anti-Flag M1 Agarose gel. Beads were than washed 3× with 10 mM Tris pH 8.0 100 mM NaCL containing 0.05% saponin and 2× with 10 mM Tris pH 8.0 100 mM NaCL without saponin. ^FT^TTR^A25T^ was eluted at 4 degrees C overnight with 50 μL of 0.1 M triethylamine dissolved in water pH 11.5 with gentle shaking. Eluted TTR molecular weights were analyzed by LC-MS, as previously described [33].

### Statistical Methods

All p-values were calculated using a paired student’s t-test.

## ACKNOWLEDGEMENTS

We thank Evan Powers, Bibiana Rius, and Jessica Rosarda for critical reading of this manuscript. We thank Jeff Kelly (TSRI) for access to the uPLC instrument and compound **1**, Taf, and Taf-S. This work was funded by NIH NS092829 to RLW.

## AUTHOR CONTRIBUTIONS

ICR and RLW conceived of the experiments, performed the experiments, analyzed the results, and wrote the manuscript.

---

**Expanded View Figure 1.**
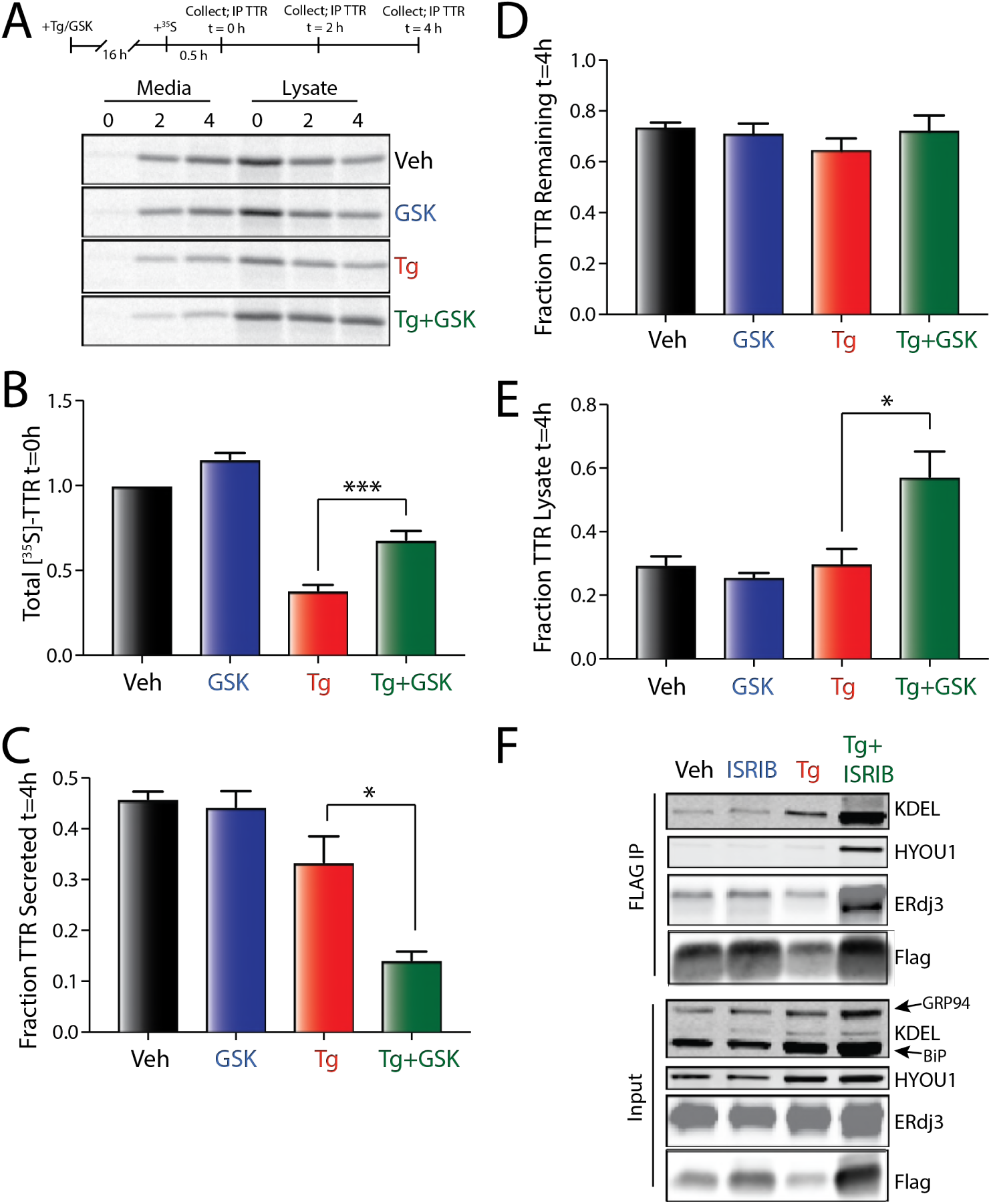
(Relevant to Figure 1) **A.** Representative autoradiogram of [^35^S]-labeled ^FT^TTR^A25T^ in lysates and media collected from HEK293T cells pretreated for 16 h with thapsigargin (Tg; 500 nM) and/or GSK (300 nM). The protocol for the experiment is shown above. **B.** Graph showing the total amount of [^35^S]-labeled ^FT^TTR^A25T^ at t = 0 h in HEK293T cells pretreated for 16 h with thapsigargin (Tg; 500 nM) and/or GSK (300 nM). Data are shown normalized to vehicle-treated cells. Error bars show SEM for n=3 independent experiments. A representative autoradiogram is shown in **Fig. EV1A.** **C.** Graph showing the fraction [^35^S]-labeled ^FT^TTR^A25T^ secreted at t = 4 h in HEK293T cells pretreated for 16 h with thapsigargin (Tg; 500 nM) and/or GSK (300 nM). Fraction secreted was calculated as described in **Materials and Methods**. Error bars show SEM for n=3 independent experiments. A representative autoradiogram is shown in **Fig. EV1A.** **D.** Graph showing the fraction [^35^S]-labeled ^FT^TTR^A25T^ remaining at t = 4 h in HEK293T cells pretreated for 16 h with thapsigargin (Tg; 500 nM) and/or GSK (300 nM). Fraction remaining was calculated as described in **Materials and Methods**. Error bars show SEM for n=3 independent experiments. A representative autoradiogram is shown in **Fig. EV1A.** **E.** Graph showing the fraction [^35^S]-labeled ^FT^TTR^A25T^ in lysates at t = 4 h in HEK293T cells pretreated for 16 h with thapsigargin (Tg; 500 nM) and/or GSK (300 nM). Fraction lysate was calculated as described in **Materials and Methods**. Error bars show SEM for n=3 independent experiments. A representative autoradiogram is shown in **Fig. EV1A.** **F.** Representative immunoblot showing anti-FLAG IPs and inputs from lysates prepared from HEK293T cells expressing ^FT^TTR^A25T^ and pretreated for 16 h with Tg (500 nM) and/or ISRIB (200 nM) *indicates p<0.05; **indicates p<0.01 for a paired two-tailed t-test.

**Expanded View Figure 2.**
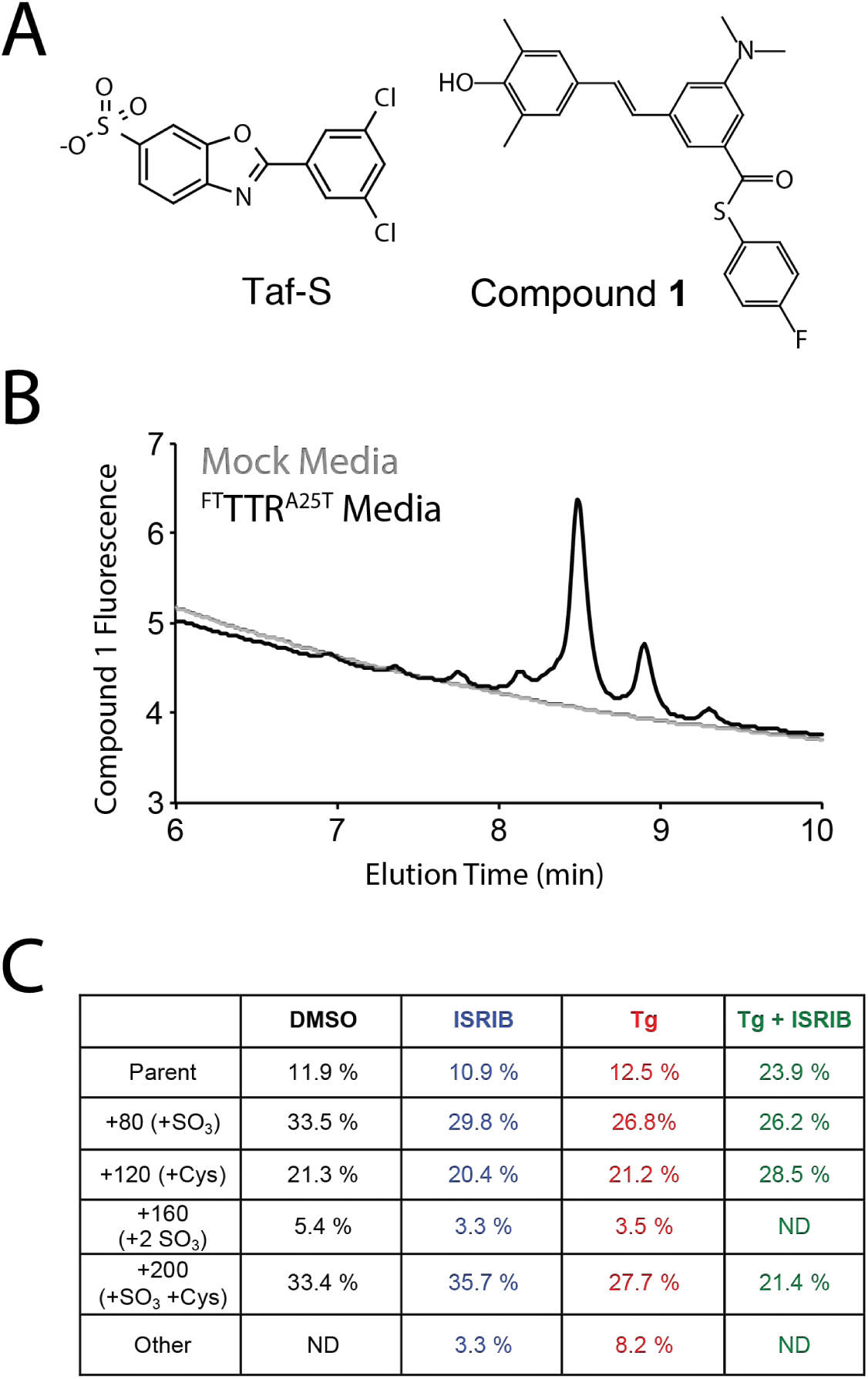
(Relevant to Figure 2) **A.** Structures of tafamidis-sulfonate (Taf-S) and the fluorogenic TTR ligand compound **1.** **B.** Chromatogram of conditioned media prepared in the presence of Taf-S (10 μM) on mock-transfected HEK293T cells or HEK293T cells expressing ^FT^TTR^A25T^. ^FT^TTR^A25T^ tetramers were separated on anion exchange chromatography using the UPLC system and visualized by compound **1** fluorescence. **C.** Table showing the relative populations of unmodified and posttranslationally modified ^FT^TTR^A25T^ in conditioned media prepared on HEK293T cells treated for 16 h with Tg (500 nM) and/or ISRIB (200 nM).

**Expanded View Figure 3.**
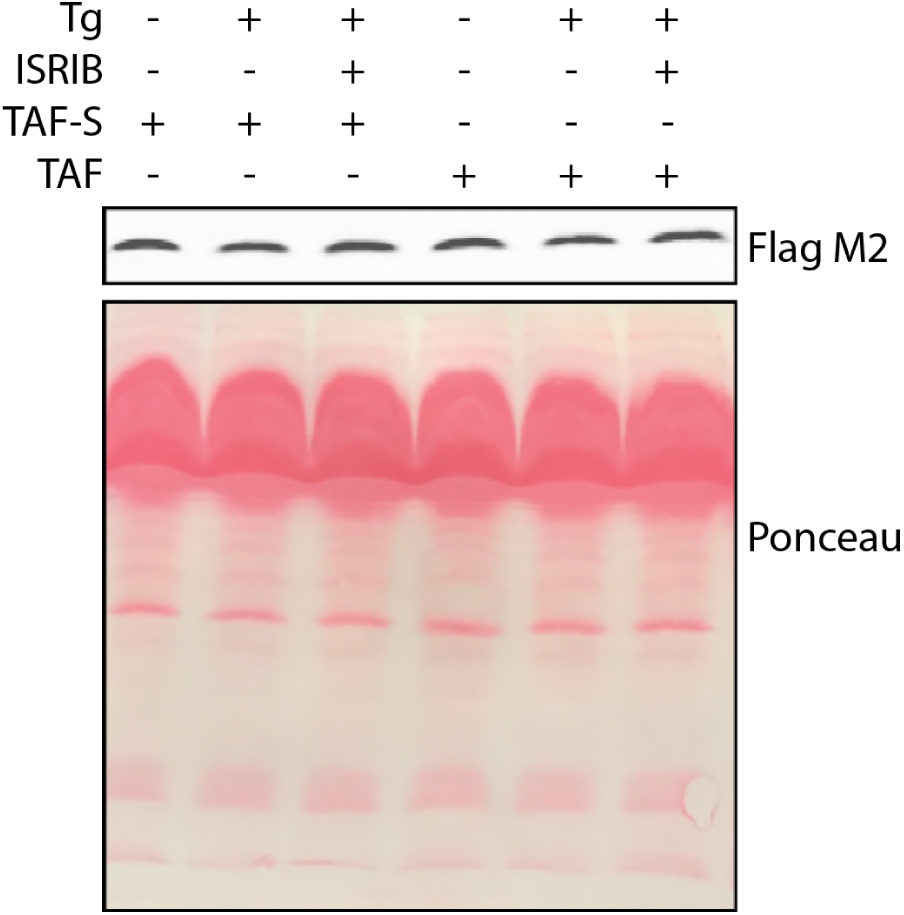
(Relevant to Figure 3) Representative immunoblot of ^FT^TTR^A25T^ in conditioned media prepared in the presence of Taf-S (10 μM) or Taf (10 μM) on HEK293T cells treated for 16 h with Tg (500 nM) and/or ISRIB (200 nM). A ponceau stain of the immunoblot is shown as a control.

**Expanded View Figure 4.**
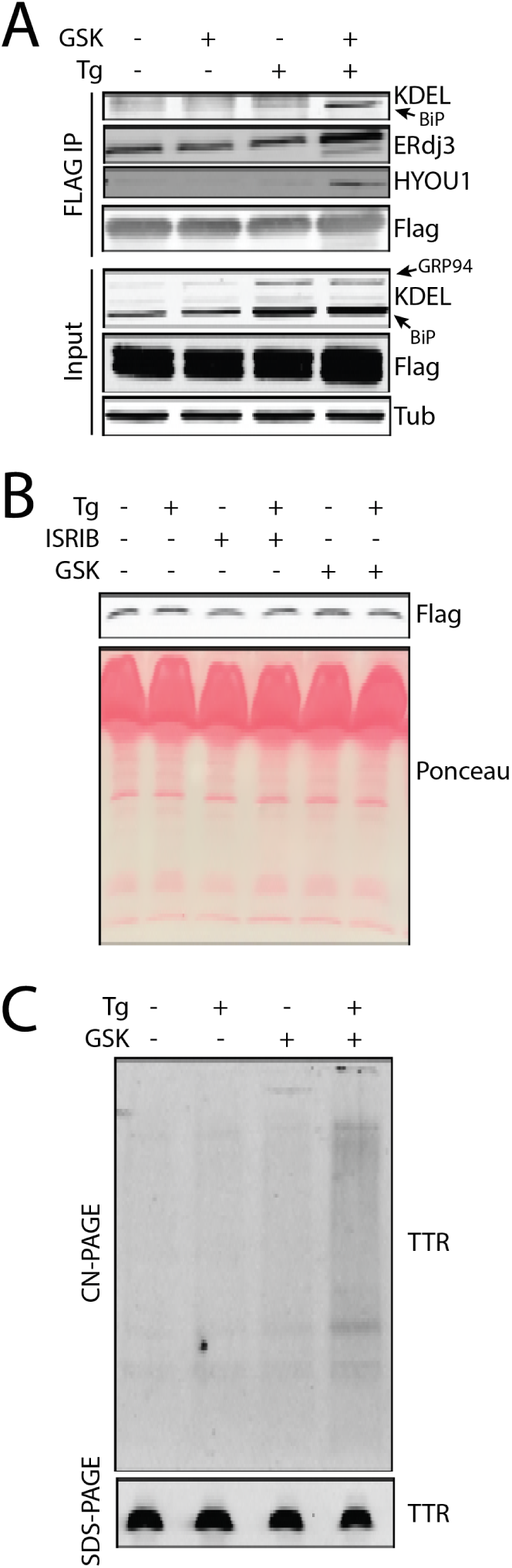
(Relevant to Figure 4). **A.** Representative immunoblot showing anti-FLAG IPs and inputs from lysates prepared from HEK293Tcells expressing ^FT^TTR^WT^ and pretreated for 16 h with Tg (500 nM) and/or GSK (300 nM). **B.** Representative immunoblot of ^FT^TTR^WT^ in conditioned media prepared in the presence of Taf-S (10 μM) on HEK293T cells treated for 16 h with Tg (500 nM), ISRIB (200 nM), or GSK (300 nM). A ponceau stain of the immunoblot is shown as a control. **C.** Representative CN-PAGE/immunoblot of ^FT^TTR^WT^ aggregates in conditioned media prepared in the presence of Taf-S (10 μM) on HEK293T cells treated for 16 h with Tg (500 nM) and/or GSK (300 nM). A SDS-PAGE/immunoblot showing total ^FT^TTR^WT^ is shown as a control.

